# Genetic Modifiers of Cognitive Function: Novel Genetic Associations with Memory, Executive Function, and Attention in the Long Life Family Study

**DOI:** 10.64898/2026.02.02.703284

**Authors:** Hannah J. Lords, Mengze Li, Zeyuan Song, Anastasia Leshchyk, Harold Bae, Nicole Roth, Megan Short, Thomas T. Perls, Moonil Kang, Ting Fang A. Ang, Stephanie Cosentino, Sandra Barral, Marianne Nygaard, Konstantin Arbeev, Stacy L. Andersen, Anastasia Gurinovich, Paola Sebastiani

## Abstract

Cognitive performance is central to health and quality of life. Studying the factors that sustain performance may offer insights into maintaining cognitive function into advanced ages and may inform strategies to promote healthy cognitive aging. To identify single nucleotide polymorphisms (SNPs) underlying cognitive function, we performed genome-wide association studies (GWAS) of nine neuropsychological test scores capturing performance in three cognitive domains in 2,455 participants of the Long Life Family Study (LLFS). We identified 12 variants in seven tests and three domains that reached genome-wide significance (p < 5×10⁻⁸). Three rare (minor allele frequency, European population MAF<0.01) protective, intronic variants, rs190287985 (*CCSER1*), rs75730801 (*FHOD3*), and rs552842447 (*LINC00508*), were uniquely associated with semantic fluency, phonemic fluency and number span forward, respectively. Two rare deleterious variants, rs556333682 and rs188304645, were associated with performance on the Hopkins Verbal Learning Test-Revised (HVLT-R) learning trials and lie in ischemia-related genes *SH3TC1* and *RPH3A*. Another variant, rs180691759, associated with HVLT-R delayed recall, was proximal to *RSPO3* and linked to decreased *ECHDC1* expression, implicating ischemic and unexplained Ethylmalonic acid (EMA) pathways. We compared the results with GWASs of a general cognitive factor (GCF) and reaction time (RT) in the UK Biobank and meta-analysis results of the UK Biobank, CHARGE and COGENT by Davies et al. We found that rs535509651 associates with HVLT-R delayed recall in the LLFS and nominally associates (p<0.05) with GCF, and that rs10424537 associates with Immediate Logical Memory in the LLFS and nominally associates with RT. Furthermore, we identified 5 genome-wide significant loci in Davies et al. that reached loci adjusted significance in the LLFS (GCF p<0.05/128 and RT p<0.05/39). Each locus was associated with a single cognitive domain in the LLFS. We annotated the genome-wide significant results with quantitative trait loci (QTL) analyses of whole blood transcriptomic, serum metabolomic data, and gene set enrichment analyses (GSEA) using all nominally significant transcripts (p < 0.05/12). QTL analyses discovered 1 SNP-transcript (*ECHDC1*, p < 3 × 10^−6^), no SNP–lipid (p < 2 × 10^−4^), and no SNP–polar metabolite (p < 2 × 10^−4^) associations. Gene set enrichment analyses identified three pathways at FDR < 0.05. Our findings provide insight into the domain-specific genetic architecture of cognitive function.

## 1 Introduction

Modern improvements to health care have led to more people surviving at older ages. However, by the age of 70, approximately two-thirds of Americans will experience some form of cognitive impairment (*1*). Even if an individual avoids dementia or mild cognitive impairment, normal cognitive aging can impact everyday well-being and quality of life (*2, 3*). Moreover, strain from other health issues may deleteriously impact cognitive function (*4*). With an ever-aging population and with it an ever-increasing strain on cognitive function, there is growing pressure to identify protective measures and develop therapies to promote healthy cognitive aging.

Cognitive function is a complex trait. Within a single individual, one can maintain one aspect (ex. speed), improve in another (ex. verbal fluency), and deteriorate in a third (ex. memory) (*2, 5–7*). Cognitive function can be measured via neuropsychological batteries, consisting of multiple tests that measure various aspects of cognitive function (*8, 9*). Multiple studies indicate that there are genetic influences acting on cognitive function (*8, 10–14*). The fact that these influences vary depending on the aspect of cognitive function measured (*8, 14–16*) and the age group of the individuals studied (*14, 17*), in conjunction with the differential neuropsychological profiles observed across a range of clinical conditions renders it unlikely that all aspects of cognitive function are implemented through the same mechanisms. These observations motivate studies of individual cognitive functions and favor this approach over combining the tests into a general cognitive function score.

Analyses of long-lived individuals and their relatives show delayed onset and compression of morbidity for a variety of diseases (*18–20*). Several investigations of participants in the Long Life Family Study (LLFS) in comparison to various control groups suggest that LLFS participants perform better on a variety of cognitive tests (*21–23*). Cosentino et al. (*13*) reported a reduced prevalence of cognitive impairment in the LLFS offspring generation compared to spousal controls. Andersen et al (*19*) found that supercentenarians (age 110–119 years), semi-supercentenarians (age 105–109) and centenarians (age 100–104) had delayed cognitive impairment compared to nonagenarian controls (age 87–99). Sebastiani et al. (*20*) reported that LLFS participants experience delayed onset of dementia compared to New England Centenarian Study centenarians and controls. Beker et al. (*24*) reported that some centenarians (age > 100) maintain high cognitive performance despite exposure to the number one risk factor for Alzheimer’s disease (i.e., advanced age), suggesting that a subset of centenarians have mechanisms of cognitive resilience. Because of this better performance on cognitive tests, delayed onset of cognitive impairment and the fact that these traits are heritable (*10, 25*), long-lived individuals and their families are ideal for studies aiming to identify cognitive protective mechanisms as they are likely enriched for protective variants. This theory is supported by the growing number of longevity studies producing associations suggestive of more youthful molecular profiles or protective mechanisms (*10, 26, 27*). Furthermore, while large-scale GWASs of cognition have identified associated variants, the total explained variance is much lower than estimated heritability(*14*). Bearden and Glahn (*14*) suggest rare variants may be critical for understanding the genetic architecture of cognition in the normative range. As a family-based study, the LLFS is uniquely positioned to identify rare variants associated with cognitive performance.

Here, we perform genome-wide association analyses of cognitive function in the LLFS, a study of families with multiple long-lived individuals. We leverage a battery of nine neuropsychological tests and the expected enrichment of longevity specific rare variants aiming to identify novel variants associated with specific cognitive functions. We supplement our analyses with associations between our detected variants and transcripts, lipids, and polar metabolites. Finally, we annotate a previous high-powered study of general cognitive function in the UK biobank and a meta-analysis of general cognition in the UK Biobank, CHARGE and COGENT participants by Davies et al. (*12*) with the underlying cognitive domains to create a more nuanced view of what the variants are capturing.

## 2 Results

### 2.1 Demographics characteristics

Table 1 summarizes the demographic characteristics of participants enrolled in the LLFS. The LLFS enrolled individuals from long-lived families between 2006 and 2009 and collected biological samples and extensive phenotypic data. A second visit was conducted between 2013 and 2018 and generated a more comprehensive assessment of cognitive function adding three neuropsychological tests to the six available in the first visit (*25*). We used the data of the second in-person assessment to identify single nucleotide polymorphisms (SNPs) associated with nine neuropsychological tests. A description of each test is in Supplementary Table 1. For a description of the test-specific data availability, phenotype statistics and covariate distributions see Supplementary Tables 1 and 2.

**Table 1:** Summary of LLFS Data.

| **Analysis** | **Participants**<br>**# Samples** | **Sex**<br>**Female(%)** | **Age**<br>**mean(sd)** | **Education**<br>**mean(sd)** | **Field Center**<br>**Denmark(%)** | **Date of Birth**<br>**After 1935(%)** |
| --- | --- | --- | --- | --- | --- | --- |
| GWAS | 2455 | 54.6% | 73(10) | 12(3) | 30.9% | 79.1% |
| Lipid QTL | 3572 | 56.5% | 72(20) | 12(4) | 22.5% | 61.1% |
| Polar QTL | 3573 | 56.5% | 72(20) | 12(4) | 22.5% | 61.1% |
| Expression QTL | 1628 | 56.1% | 70(20) | 11(4) | 34.6% | 63.9% |

Analysis: analysis conducted. Participants # Samples: Sample size. Sex: percent female. Age: the mean (sd) age of the samples. Education: the mean (sd) years education. Field Center: percent of participants from the field center in Denmark. Date of Birth After 1935: the percent of participants born after 1935.

**Table 2:** Genome-wide Significant Results.

| Phenotype | Chr | Pos | rsID | Effect Allele | REF Allele | Est | Est.SE | Est Percent of Score Range | P-value | EAF | Eur AF | Nearest Gene(s) | Genomic Function |
| --- | --- | --- | --- | --- | --- | --- | --- | --- | --- | --- | --- | --- | --- |
| hvltotcor4 | 1 | 40,307,373 | rs187863974 | T | C | -5.930 | 1.00 | -49.4% | 1.53e-08 | 0.002 | 0.000 | COL9A2 | intronic |
| hvltotrecall | 4 | 8,241,275 | rs556333682 | A | G | -12.700 | 2.30 | -35.3% | 1.87e-08 | 0.001 | 0.000 | SH3TC1 | downstream |
| animal | 4 | 90,705,099 | rs190287985 | G | T | 15.700 | 2.70 | 34.9% | 5.09e-09 | 0.001 | 0.002 | CCSER1 | intronic |
| hvltotcor4_no0 | 5 | 59,820,828 | rs142162066 | C | G | -2.390 | 0.43 | -21.7% | 2.73e-08 | 0.007 | 0.012 | PDE4D | intronic |
| hvltotcor4 | 6 | 126,833,110 | rs180691759 | G | A | -6.300 | 1.00 | -52.5% | 1.66e-09 | 0.002 | 0.001 | MIR588;RSPO3 | intergenic |
| hvltotcor4 | 10 | 27,445,277 | rs535509651 | C | A | -4.600 | 0.82 | -38.3% | 2.10e-08 | 0.003 | 0.004 | PTCHD3;RAB18 | intergenic |
| hvltotrecall | 12 | 112,709,168 | rs188304645 | G | A | -5.680 | 1.00 | -15.8% | 4.41e-08 | 0.005 | 0.003 | RPH3A | intronic |
| digitfwd | 12 | 127,957,990 | rs75730801 | T | A | 1.440 | 0.26 | 13.1% | 2.11e-08 | 0.013 | 0.005 | LINC00508 | ncRNA_intronic |
| fastot | 18 | 36,647,892 | rs552842447 | C | G | 13.900 | 2.30 | 15.3% | 3.11e-09 | 0.007 | 0.006 | FHOD3 | intronic |
| logmem_imd | 19 | 55,255,621 | rs10424537 | C | T | -0.634 | 0.11 | -2.5% | 1.90e-08 | 0.482 | 0.462 | PPP6R1 | intronic |
| hvltotcor4_no0 | 20 | 47,913,267 | rs183261378 | T | C | -5.260 | 0.91 | -47.8% | 6.27e-09 | 0.002 | 0.001 | SULF2;LINC01522 | intergenic |
| hvltotrecall | 22 | 12,531,750 | rs1263369249 | T | C | -8.700 | 1.60 | -24.2% | 2.98e-08 | 0.002 | 0.000 | LOC102723769;NONE | intergenic |

Chr: the chromosome. Pos: base pair position in GRCh38. rsID: reference SNP ID from dbSNP. Effect Allele: code allele. REF Allele: reference allele from GRCh38. Est: estimated change in the test score per effect allele. Est.SE: the standard error of Est. Est Percent of Score Range: The estimated effect as a percentage of the range of the test scores. EAF: frequency of the effect. Eur AF: effect allele frequency in gnomADv41 (non-Finish European). Nearest Gene(s): the upstream and downstream closest reference genes to the SNP.

### 2.2 Genome-wide association studies of nine neuropsychological tests identify novel genetic modifiers

We first estimated the SNP-based heritability of the nine neuropsychological tests and pairwise genetic correlations (r_g_). We found that the nine neuropsychological tests had heritabilities ranging between 17% and 45% (Figure 1, Supplementary Table 3). Wojczynski et al. (*25*) also performed pedigree-based heritability analyses for Number Span Forward, the Digit Symbol Substitution Test, Semantic Fluency, HVLT-R learning trials, Number Span Backward, Logical Memory immediate recall, and Logical Memory delayed recall in the LLFS and estimated higher heritabilities for all these tests compared to our estimates. Despite the order of magnitude, the ranking of heritability estimates in the two analyses is consistent except for the Number Span Backward and HVLT-R learning trials (Supplementary Table 3). The heritability of learning and memory tests was low (17-20%) compared to executive function (23-45%) and attention (37%) tests (Figure 1, Supplementary Table 3). The genetic correlation was as low as 0.05 Digit Span Backwards and Logical Memory delayed recall, suggesting that the various tests had unique genetic architectures (Figure 1, Supplementary Table 4).

**Figure 1:**
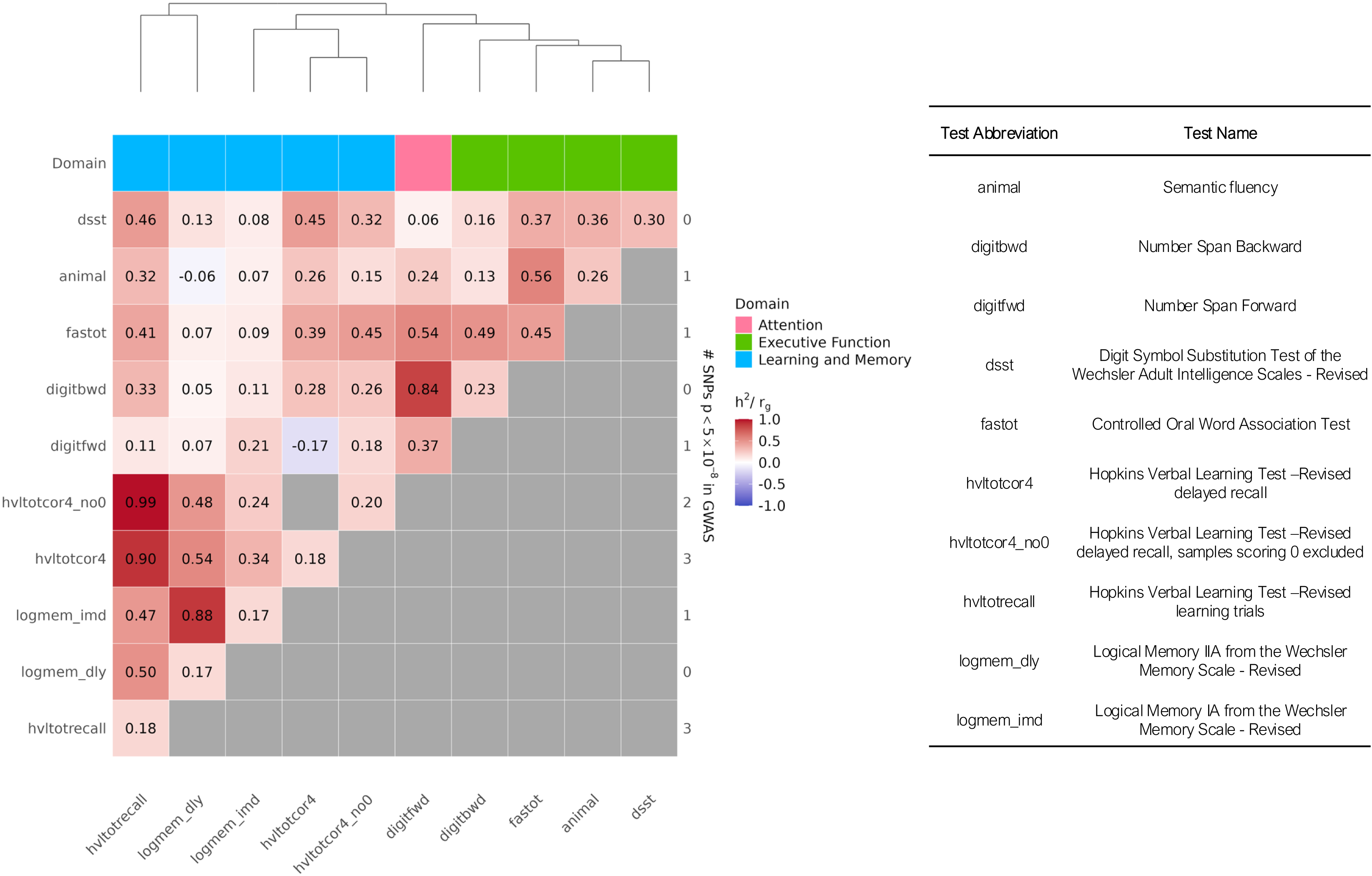
Heritability and Genetic Correlation of Neuropsychological Tests. y-axis and x-axis labels display test abbreviations. The right-hand y-axis indicates the number of genome-wide significant SNPs (p < 5×10⁻⁸) identified in each corresponding GWAS. The Domain row denotes the cognitive domain best captured by each test. The legend maps the test abbreviations used in the plot to the test name.

Next, we conducted Genome-wide association studies (GWASs) of the nine neuropsychological test scores for 9 million SNPs (Supplementary table 1). Across all nine cognitive metrics GWASs, we identified 12 genome-wide significant (p<5×10^−8^) associations in seven neuropsychological tests scores. No SNP was associated with multiple tests scores. Among these SNPs, 10 were rare variants (effect allele frequency < 0.01 in the GNOMAD4.1 European population) and three of the rare variants were associated with better performance. The only genome-wide significant SNP that was sensitive to APOE was the association between rs552842447 and phonemic fluency [fastot] (Supplementary Table 5).

Two SNPs reached genome-wide significant association with semantic fluency [animal] and phonemic fluency [fastot] in the executive function domain. We estimated that individuals with the effect allele, G, for rs190287985 (p = 5.09×10^−9^) on average have an improvement in semantic fluency performance of 15.73 points per effect allele, an improvement of 34.9% of the test score range (SR). This SNP is an intronic variant in *CCSER1*. For the phonemic fluency variant, rs552842447 (p=3.11×10^−9^), we calculated that on average individuals with the effect allele, C, have an improvement of 13.89 points per effect allele, an increase of 15.3% SR. This variant is intronic to *FHOD3*. We identify one variant, rs75730801 with effect allele T (p=2.11×10^−8^), associated with the attention domain Number Span Forward test [digitfwd]. This variant is a beneficial variant (est=1.44, 13.1% SR) located within the intronic region of the non-coding RNA *LINC00508.* It is rare in the general European population (EAF = 0.005), but not in the LLFS (EAF = 0.013) (Table 2, Figure 2).

**Figure 2:**
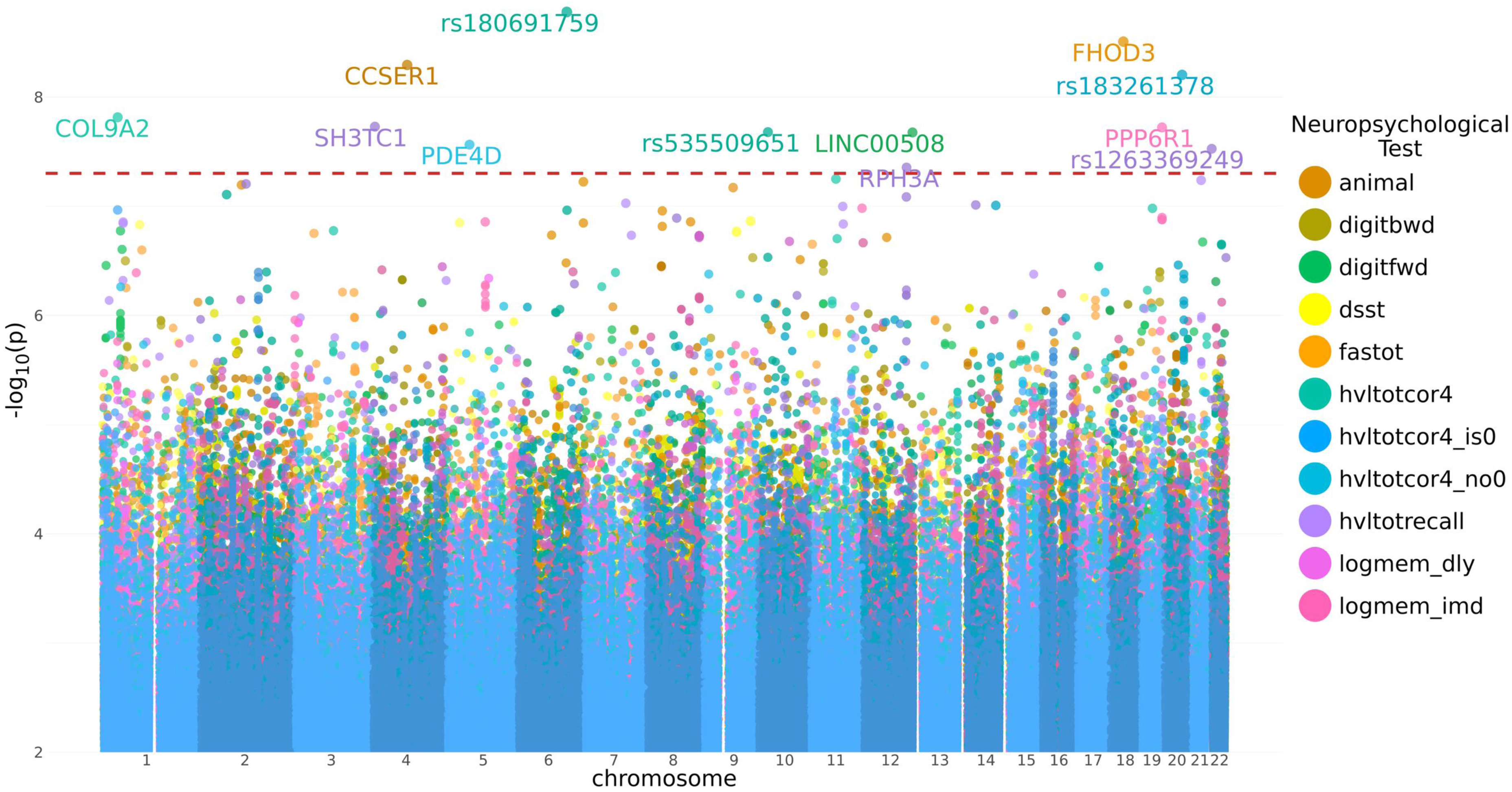
Manhattan Plot of All Neuropsychological Test GWASs. Overlapping color coded Manhattan plot for each GWAS. The dotted red line indicates genome wide significance (p<5×10^−8^). Each genome wide significant point is labeled with either the rsID if the SNP is intergenic or the Gene name if it is within any component of a gene.

We identified nine SNPs associated with tests in the learning and memory domain (p < 5×10^−8^): four intronic, one within the 3’UTR, and four intergenic. Seven of these variants were rare. All significant variants had deleterious estimated effects for the minor allele (Table 2, Figure 2). We identified three SNPs associated with HVLT-R delayed recall [hvltotcor4], two SNPs associated with HVLT-R delayed recall when samples scoring 0 were excluded [hvltotcor4_no0], and three SNPs associated with performance on the HVLT-R learning trials [hvltotrecall]. We estimate that individuals with these variants will perform worse by more than 15% SR. The effect allele of rs142162066, associated with hvltotcor4_no0 (est=-2.39, –21.7%SR, p=2.73×10^−8^) is not rare in the general European population (EAF = 0.012), but is rare (EAF = 0.007) in the LLFS. The final variant (rs10424537, effect allele = C) was associated with Logical Memory immediate recall (est=-0.63, –2.5% SR, p=1.90×10^−8^). With an effect allele frequency of 0.482 in the LLFS, rs10424537 is a common variant. As seen in Figure 2, the Logical Memory immediate recall hit is a singleton. This aligns with the linkage structure of the region (Supplementary Figure 1).

### 2.3 Comparison with Published Results

To replicate our results, we compared all our genome-wide significant results to the Davies et al. (*12*) GWAS results of general cognition factor (GCF) and/or reaction time (RT) estimates. For this replication, we limited comparison to the results generated from UK Biobank participants as only their results were publicly available. Only nine of our genome-wide significant SNPs had data available for comparison (Table 3, Figure 3a). Of these, one locus of the nine was associated with GCF (GCF p <0.05 / number of LLFS hits for that test), and another was associated with RT (RT p < 0.05 / number of LLFS hits for that test). The SNP, rs535509651, was previously reported by Davies et al. (*12*) to be associated with GCF (p=9.67×10^−3^, z=-2.59). In our analysis, we observed a significant association with HVLT-R delayed recall [hvltotcor4] (p=2.10×10^−8^, est=-4.60, –38.3% SR). Given that the directions of both effects are deleterious, the finding reported by Davies et al is replicated (Table 3, Figure 3a). On the other hand, the SNP rs10424537 was previously reported by Davies et al. (*12*) to have a protective effect on reaction time (p=2.74×10^−3^, est=-0.0038). In our analysis, however, we observed a deleterious association with Logical Memory immediate recall [logmem_imd] (p=1.90×10^−8^, est=-0.63, –2.5% SR). Given the lack of concordance in effect direction, we do not consider this locus to represent a replication (Table 3, Figure 3b).

**Figure 3:**
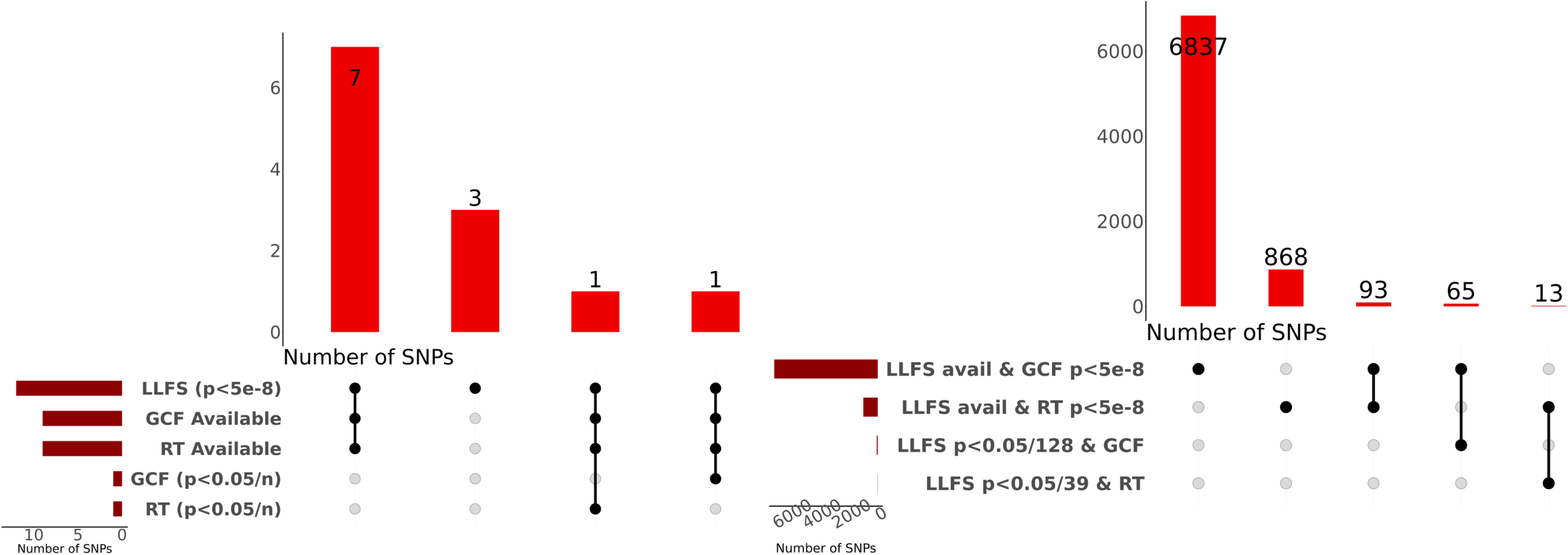
Upset Plots of Comparison with the Davies et. al Results. Upset plots of the SNPs available in the LLFS and Davies et. al GWASs broken down by significance. **Figure 3a** shows the 12 genome-wide significant SNPs, seven were available but did not reach adjusted significance in Davies et. al. Three were only available in the LLFS. One was significant in the LLFS and Davies et. al general cognition factor when n is the number of hits significant for the associated test. One was significant in the LLFS and adjusted significant for Davies et. al reaction time (for more details see replication methods section 5.2). **Figure3b** shows the 67 general cognition factor and 13 reaction time results that were significant (p<5×10^−8^) in Davies et. al and reached adjusted significance in the LLFS based on the number of loci identified in the corresponding Davies et al GWAS (for more details see replication methods section 5.2).

**Table 3:**
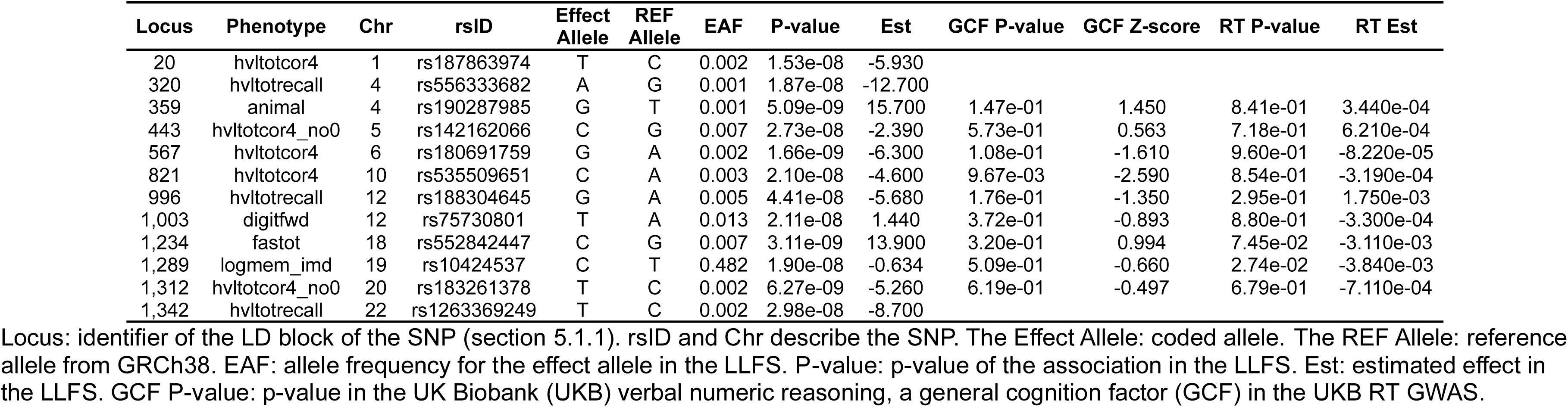
LLFS p<5×10-8 SNPs with Davies et.al results.

In addition to comparing our significant results to Davies et al. (*12*), we also compared Davies et al.’s significant results, from the CHARGE, COGENT and UK Biobank cohort meta-analysis, to all our LLFS cognitive GWAS results, regardless of p-value. Davies et al. (*12*) identified 11,600 SNPs in 148 loci that were associated with their GCF variable. Of these, 6,995 SNPs in 128 loci were available in LLFS and 65 SNPs in four loci were significantly associated with an LLFS cognitive metric at p < 0.05/128 (Figure 3b). Locus 138 consists of two SNPs; one SNP was associated only with Logical Memory delayed recall [logmem_dly] and one associated with both immediate [logmem_imd] and delayed Logical Memory. Thus, this Locus appears to be specific to Logical Memory. Locus 246 contains 51 SNPs associated with the failure to earn even a single point on the HVLT-R delayed recall test [hvltotcor4_no0]. Locus 454 includes 11 SNPs; 10 SNPs were associated with delayed Logical Memory [logmem_dly], while HVLT-R delayed recall (regardless of inclusion of samples scoring zero) [hvltotcor4, hvltotcor4_no0] associated with one SNP. The final locus contains a single SNP which was associated with DSST. All the results are summarized in Supplementary Table 6 and Supplementary Figure 2. Davies et al. also identified 2,022 SNPs in 42 loci associated with reaction time. Of these, 974 SNPs in 39 loci were available in LLFS and 13 SNPs in locus 157 were significantly associated (p < 0.05/39) with performance on the Digit Span Forward test [digitfwd] (Figure 3b). These results are summarized in Supplementary Table 7. Thus, each locus seems to capture a specific cognitive domain, and some loci seem to be capturing even more specific aspects of cognitive function.

### 2.4 Quantitative Trait Loci (QTL) Analyses

We correlated gene expression in blood, and serum lipidomics and polar metabolomics with the 12 genome-wide significant SNPs using data from the first in-person visit, which has the most samples available (Table 1). We identified one association between our genome-wide hits and transcript expression level [linear association] (p<0.05/16,301), and no associations between our genome-wide hits and whether the transcript was expressed [logistic association] (p<0.05/1,102). Only ethylmalonyl-CoA decarboxylase 1 (*ECHDC1*) transcript expression was significantly associated with the SNP *rs180691759* (p=1.06×10^−8^). The analysis showed that the expression of *ECHDC1* decreased by e^−0.7388^+1 for each copy of the G allele, which we found to be associated with worse performance in HVLT-R delayed recall [hvltotcor4] (Supplement Table 9). We did not identify any lipid (p < 0.05/188) or polar (p < 0.05/220) metabolites that were associated with our significant GWAS hits (Supplement Table 9).

Additionally, we performed gene set enrichment for the SNPs that reached genome-wide significance in the LLFS and the Hallmark (N=3,361 genes) and Kyoto Encyclopedia of Genes and Genomes (KEGG; N=3,550) compendia of genes with blood transcript expression data available in the LLFS. When considering only the linearly associated transcripts, only the logistic associated transcripts or both linear and logistic combined suggestive significant (p<0.05/12) SNP associated transcripts, we did not identify any significantly enriched (q < 0.05, using Benjamini-Hochberg False Discovery Rate correction) gene sets in the Hallmark compendium (Supplementary Table 9). For the KEGG compendium, the rs1263369249 effect allele is associated linearly with 19 genes in the ribosome pathway. HVLT-R learning trial associated SNPs were associated with 23 genes in the ribosome pathway. The rs188304645 effect allele was associated linearly with three genes in the KEGG JAK-STAT signaling pathway. Finally, the HVLT-R learning trials [hvltotrecall] associated SNPs were associated with 11 genes in the KEGG oxidative phosphorylation pathway. This enrichment is specific to the HVLT-R learning trials associated SNPs and was not identified for any other test or domain (Supplementary Table 11).

## 3 Discussion

We estimated heritability and genetic correlation of nine neuropsychological test scores. Next, we conducted GWASs of these test scores. We discovered 12 SNPs that reached genome-wide significance, and generated omics signatures for these SNPs. We compared our results to the results reported in Davies et al.

(*12*). The heritability and the genetic correlation vary by test. We identified four GCF and one RT loci associated with specific domains. We identified one deleterious common variant (EAF > 0.05), three protective uncommon (EAF<0.05) variants and eight detrimental uncommon variants, including one linked to *ECHDC1* expression. The SNPs associated with the HVLT-R learning trials were nominally associated with transcripts of 11 genes in KEGG’s oxidative phosphorylation pathway.

### The learning and memory domain might depend on a smaller number of SNPs that are more easily detected than other domains

Our heritability analyses showed that neuropsychological tests related to executive function and attention are more heritable than those assessing learning and memory (Figure 1, Supplementary Table 3). We found that heritabilities from Wojczynski et al. (*25*) were systematically larger than those reported in this study (Supplementary Table 3). However, SNP-based heritability often returns lower estimates than pedigree-based heritability (*14*). Furthermore, in our analyses, we adjusted the model by education, which is a well-known confounder of cognitive performance, where the estimates of Wojczynski et al. (*25*) did not adjust by education. Despite these discrepancies, Wojczynski et al. (*25*) replicated the pattern of learning and memory domain tests having lower heritability except for Number Span Backward [digitbwd] being estimated as slightly more heritable than HVLT-R learning trials [hvltotrecall]. This pattern of heritability suggests that performance on executive function and attention tasks is influenced more strongly by genetic or familial factors. However, our genome-wide association analyses identified more variants associated with learning and memory than executive function or attention (Figure 1, Table 2). Furthermore, we found that the GCF which essentially combines the effects across domains, captures learning and memory affecting SNPs better than any other domain. We identified four loci that reached genome-wide significance in the GCF and the LLFS. Three of these loci were specific to learning and memory tests (Supplementary Table 6), suggesting that the learning and memory SNPs are easier to detect, even when masked by the transformation into general cognition. These results suggest a difference in the genetic architectures of the domains and we hypothesize that learning and memory processes may rely on a smaller number of genetic variants with larger, more independent effects, whereas executive function and attention may be governed by a more polygenic architecture with many variants of individually small effect or perhaps with effects that are dependent on having a specific combination of variants or circumstances.

### Cognitive domains have distinct genetic architectures motivating test specific analyses

Consistent with the low genetic correlation between traits (average r_g_=0.32), we observed no overlap in the variants associated with different tests (Table 2). This absence of shared loci supports the presence of domain-specific genetic influences on individual cognitive domains (Figures 1 and 2). Replication analyses using GWAS results from Davies et al. (*12*) provided further insight. Reaction time variants from the Davies et al. (*12*) study corresponded to our Digit Span Forward test [digitfwd] (Supplementary Table 7), consistent with a domain specific genetic architecture. SNPs associated with the general cognition factor from Davies et al. also showed concordance in a test specific manner, most prominently with learning and memory tests (Supplement Table 6). We identified four loci that reached adjusted significance in the LLFS, and genome-wide significance in Davies et al. (*12*). Each of these loci, associates with a specific domain or test in the LLFS. Locus 246 contains 51 SNPs concordantly associated with the GCF in Davies et al. (*12*) and with the failure to earn even a single point on the HVLT-R delayed recall test [hvltotcor4_is0] in the LLFS. Likewise, GCF locus 859 only associated with the Digit Symbol Substitution Test [dsst]. GCF locus 138 is specific to the Logical Memory tests [logmem_imd, logmem_dly], while GCF locus 454 is specific to the delayed recall tests [logmem_dly, hvlotcor4, hvltotcor4_no0] (Supplementary Tables 6, Supplementary Figure 2B). These results underscore the value of analyzing cognitive phenotypes separately rather than aggregating them into a general cognitive factor, as such aggregation may obscure important domain specific signals or lose functional nuance. These results are consistent with growing evidence of the importance of investigating factors that affect specific cognitive domains, though for the purposes of reducing measurement error and minimizing Type I error, composites of domain group tests appear to be the most effective (*28–31*).

### Oxidative stress is a potential mechanism of cognitive vulnerability

Several of the identified learning and memory-associated variants lie near genes involved in the brain’s response to ischemia and oxidative stress, pointing toward a potential mechanism of cognitive vulnerability. The SNP rs556333682 that was associated with HVLT-R learning trials [hvltotrecall] (est = –12.74, –35.3% SR, p = 1.87 × 10⁻⁸) lies downstream of SH3 Domain And Tetratricopeptide Repeat-Containing Protein, *SH3TC1*, a gene implicated in lacunar strokes and Charcot-Marie-Tooth disease (*32, 33*). Lacunar strokes are a major cause of vascular cognitive impairment and have been strongly associated with cognitive decline (*34*). Another variant, rs188304645 (hvltotrecall; est = – 5.68, –15.8% SR, p = 4.41 × 10⁻⁸), lies within Rabphilin 3A, *RPH3A*, a gene whose protein product protects neurons from ischemia-reperfusion injury (*35*). Similarly, rs180691759 (associated with HVLT-R delayed recall [hvltotcor4]; est = –6.30, –52.5% SR, p = 1.66 × 10⁻⁹) is located between microRNA-588, *MIR588* (*36*), and R-Spondin 3, *RSPO3* (*37*), both of which are involved in cellular responses to hypoxia. This variant is associated with decreased expression of ethylmalonyl-CoA decarboxylase 1, *ECHDC1*, a metabolic proofreader that detoxifies ethylmalonic acid into butyrate (*38–40*). If this function is compromised, the resulting accumulation of toxic metabolites could exacerbate oxidative damage during ischemic reperfusion (*38–41*). Our analyses associate (p=1.06×10^−8^) rs180691759:G with decreased expression of *ECHDC1* (est=-0.73). This decrease could lead to reduced production of the protein product, leading to increased ethylmalonic acid, and increased vulnerability to ischemic reperfusion injury.

These findings converge on a plausible biological narrative: many of the learning and memory-associated variants may impair neuronal resilience to oxidative or ischemic damage, leading to poorer performance on recall-based tasks. Further work looking into the role of the identified variants in poorer memory and learning is warranted and could lead to the identification of potential intervention targets.

### Limitations

Our phenotypes are designed to capture cognitive performance which is known to be related to the brain. However, we were unable to examine how our SNPs impact brain transcription and to some extent metabolomics, since the LLFS only has blood samples. Furthermore, that the SNP transcripts are associated with a change in expression in the blood does not necessarily generalize to the brain. Additionally, we cannot comment on how transcripts expressed in the blood might interact with the blood/brain barrier. The variants we identified are rare; some found in as few as four participants. We were unable to replicate most of our results. Some of this may be due to comparing the imputed data from Davies et al.(*12*) to the whole genome sequence data in the LLFS. Imputed data is best utilized to study common variants.

### Conclusions

We performed genome wide association studies of nine neuropsychological tests, identifying 12 SNPs associated with seven tests at p<5×10^−8^. One SNP, rs180691759, associated with HVLT-R delayed recall [hvltotcor4] at genome-wide significance and associated (p=1.06 × 10^−8^) with decreased expression of the *ECHDC1* transcript. We did not find any significantly associated metabolites. We replicated one SNP with the Davies et al.’s general cognitive factor and identified a possible pleiotropic effect between a SNP associated with Immediate Logical Memory [logmem_imd] and Davies et al.’s reaction time. We find that several of our SNPs (p<5×10^−8^) and their associated transcripts (p<0.05/12) are related to ischemic response. Further, we find that learning and memory as a domain is easier to detect, perhaps due to having a simpler genetic architecture compared to other cognitive domains (specifically executive function), and that this genetic architecture appears to be distinct between domains and even between tests.

### Fundings

This work was supported by the National Institutes of Health, NIA cooperative agreements U19-AG063893 and UH2/UH3-AG064704

## 4 Materials and QC Methods

### 4.1 Study Population and Genetic Data

The Long Life Family Study (LLFS) is a family-based study focused on enrolling individuals with a family history of longevity. The study enrolled 4,981 family members from 552 families between 2006 and 2009 (*25*). Whole-genome sequencing was performed with Illumina Sequencers followed by the standard alignment and quality control steps. The details of sequencing, alignment, and quality control protocols can be found in Daw et al (*42*). Due to limited power for other ancestries, our analyses limited participants to those of European ancestry. We filtered SNPs based on missing rate < 2.5%, after subsetting by participants with cognitive test data available for each test. Furthermore, individual tests applied a minor allele count (MAC) filter (MAC < 3 or MAC < 5) to remove banding from the Manhattan plots and minimize artifacts (Supplementary Table 1, Supplementary Figures 3-12).

### 4.2 Phenotypic Data

The Long Life Family Study is a longitudinal study. The first round of data collection occurred between 2006 and 2009. Only six of the nine neuropsychological tests were available at this time. For the second visit (2013–2018), three new tests were administered for the first time. Our analyses here used the performance from the second visit for all tests. The LLFS administered the following tests at the second in-person visit: Logical Memory IA (N=2,393) and IIA (N=2,306) from the Wechsler Memory Scale – Revised (WMS-R) to measure episodic memory using immediate and delayed paragraph recall. Number Span Forward (N = 2,346) & Backward (N=2,407) to measure auditory attention and working memory. Digit Symbol Substitution Test (N=2,309) of the Wechsler Adult Intelligence Scales – Revised (WAIS-R) to quantify psychomotor processing, attention and working memory. Semantic fluency (“animals”, N = 2,407) and the Controlled Oral Word Association Test (COWA, N=2,308) to measure verbal fluency. The Hopkins Verbal Learning Test –Revised (HVLT-R) learning trials (N=2,390) measures verbal learning and delayed recall (N=2,350) measures verbal episodic memory. Additionally, because the zero scores of the HVLT-R delayed recall cause the distribution to diverge from normal, we also analyzed HVLT-R delayed recall without samples scoring zero (hvltotcor4_no0, N=2,120). We also created a logistic metric that indicates whether the participant scored zero or not (hvltotcor4_is0). Samples whose tests were marked as invalid because of poor hearing or vision, environmental distractions, experimenter errors, or other physical limitations were omitted from analyses for the invalidated tests. See Supplementary Tables 1 and 2 for breakdowns of the data available and covariate distributions for each test.

### 4.3 Expression Data

The LLFS generated whole blood RNA-seq expression data for 1,855 participants and 16,301 gene-level transcripts from their first blood draw (batch 5) with complete genotype and covariate data. We performed principal component (PC) analysis of the sample transcript expression profiles. Seven samples with PC1 or PC2 values greater than 4 standard deviations (sd) away from 5% trimmed mean for their respective component were removed as outliers, leaving 1,848 samples. Some of the transcripts have a non-normal distribution, where there appears to be two separate states, an “on” state and an “off” state. The distribution of the “on” state samples are normal, but if the “off” state samples are included, there is a second peak at zero.

To account for this, we create two separate models: a linear model of the “on” state, and a logistic model of whether the sample is in the “on” or “off” state. We created a second data set for logistic regression. This data set indicated for each transcript whether the transcript was expressed (non-zero expression, ‘TRUE’) or not expressed (expression equal to zero, ‘FALSE’) for each sample. For the logistic regression data set, we excluded transcripts from the logistic regression data if either the number of TRUE values or the number of FALSE values were less than 5, as they had too few samples to meaningfully model. The resulting logistic regression data set contained 1,848 samples and 1,102 transcripts. Using the full normalized expression data set, we created our linear regression data set. In the linear regression data set, we excluded samples with zero expression for a transcript from the analysis of that transcript. The linear regression analysis data included all transcripts regardless of how much of the data was set to missing. The linear regression data set contained 1,848 samples and 16,301 transcripts.

### 4.4 Metabolite Data

We analyzed the data of 188 lipid species for 3,572 samples, and of 220 polar metabolites for 3,573 samples. All metabolomic data was profiled in plasma. Our metabolomic workflow is described in Stancliffe et al. (*43*) and Sebastiani et al. (*44*). We analyzed the lipid and polar metabolites separately and performed all QC protocols on each batch separately. We removed sample outliers based on visual inspection of PC1 and PC2. For the lipid metabolites, we removed 5 samples, and for the polar metabolites, we removed 3 samples. All metabolites were log-transformed. We set metabolites that measured more than 4 standard deviations from the metabolite’s mean to the mean of that metabolite.

## 5 Statistical Analysis

### 5.1 Genome-wide association studies (GWASs)

For the GWAS analyses, we implemented a QTL pipeline (*45, 46*) utilizing GENESIS(*47*) linear mixed effect models. To control for family relatedness, we used a genetic relationship matrix (GRM) calculated using PC-Air and PC-relate (*45, 48, 49*) We added the first four principal components (PCs) derived from the genotype data to adjust for population structure, calculated using PC-Air (*45, 48*). We calculated the GRM and PCs once for all individuals in the analysis, despite every test having a slightly different set of samples.

Additionally, we adjusted for sex, education level as an ordinal variable capturing the participant’s highest degree, a binary variable indicating whether the participant was recruited at the Denmark field center or in the United States, age, a binary variable birth cohort (categorizing individuals based on whether they were born on or before 1935, or after 1935), and the interaction between birth cohort and age. We include birth cohort to account for the bimodal age distribution from enrolling parents and their offspring. For the hvltotcor_is0 GWAS, we used a GENESIS logistic mixed effect model with the same covariates, PCs and GRM. For the logistic GWAS, we only used SNPs for which the saddle point approximation converged in our downstream analyses. Genome-wide significant SNPs were mapped into LD blocks, using the European Ancestry LD blocks defined by MacDonald et al. (*50*).

#### 5.1.1 Additional Genetic Analyses

We identified SNPs known to correlate (R^2^>0.8) with rs10424537, using LDlink’s LDproxy tool (*51*) for the Utah residents from North and West Europe population and a ±500kb window (Supplementary Figure 1). We also modeled the association between APOE and our neuropsychological tests, using GENESIS linear mixed effect models and the same covariates as our GWASs (Supplementary Table 12), and performed APOE sensitivity analyses for our genome-wide significant SNPs. We estimated heritability using the null model from each neuropsychological test’s GWAS. Variance components were estimated by restricted maximum likelihood (REML) of the null model, and 95% confidence intervals were obtained using profile likelihood via the varcompCI function in GENESIS (*47*). We calculated the genetic correlations between each neuropsychological test by fitting a bivariate mixed effect model on the samples with data available for both tests. Each model was fit using the Genome-wide Complex Trait Analysis (GCTA) software tool (*52, 53*). We adjusted by the same covariates and GRM as for the GWASs. The neuropsychological tests were standardized for use in the bivariate mixed effect models. The heatmap was clustered using the Euclidean distance and unweighted pair group method with arithmetic mean in R v4.4.0.

### 5.2 Comparisons

We downloaded the UK Biobank (UKB) GWAS results for the verbal numerical reasoning (VNR) test and reaction time (RT) from the Edinburgh Data Share Portal (https://datashare.ed.ac.uk/handle/10283/3340) on November 12, 2024 (*12*). According to Hill et al. (*54*), the UKB VNR test has high genetic correlation to general cognitive factor (GCF) score of the CHARGE and COGENT, allowing Davies et al. (*12*) to compare across cohorts. For readability, we refer to both the meta-analysis GWASs of general cognition factor in the UKB, CHARGE and COGENT datasets and the verbal numeric reasoning (UKB only) GWAS as GCF. We merged the LLFS genome-wide significant (p < 5×10^−8^) GWAS data with the UKB GWAS data, harmonizing effect alleles. Then, we defined significance as p < 0.05/n, where n is the number of genome-wide significant SNPs associated with the neuropsychological metric in the LLFS. Finally, we compared the effect directions of the UKB SNPs to the effect directions in the LLFS for the tests. We considered the effects concordant between phenotypes if the effect directions were both protective or both deleterious. If a SNP was both significant and concordant, we considered the SNP to be replicated by Davies et al. (*12*). If a SNP was significant but the effects were not in the same direction, we did not consider them to be replicated since the SNP may not be capturing the same effect. Significance without concordance may be evidence that the SNP has pleiotropic effects, but we are unable to verify this.

Additionally, using the genome-wide significant results from Davies et al.’s (*12*) Supplementary Tables 1 (GCF) and 10 (RT), we tried to replicate the findings for the meta-analysis of CHARGE, COGENT and UK Biobank from Davies et al. with the results from the LLFS, harmonizing effect alleles. We identified variants significant in Davies et al. that had GWAS results available in the LLFS. To determine whether the SNP replicated, we filtered the LLFS p-values by p < 0.05/n, where n is the number of significant loci in Davies et al. (*12*) and available in the LLFS, n = 128 for GCF and n = 39 for RT. We defined replication the same as above.

### 5.3 QTL associations and annotations

#### 5.3.1 Expression QTLs (eQTLs)

We performed linear and logistic regression analyses to identify associations between transcripts and our genome-wide significant SNPs. For our linear regression analyses, we used the same QTL pipeline, GRM and PCs as described in section 5.1. We adjusted by sex, field center, education, birth cohort, batch effect, the participants age at the time of the blood draw, the interaction between age and birth cohort, the time in years between when the participant had their blood drawn and when the blood was prepared for RNA sequencing, and the percent of the transcript data that is intergenic for that sample. For the logistic regression, where we tried to identify SNPs associated with the “on” state, we modified the QLT pipeline (*46*) to perform logistic regression with a mixed effect model implemented in GENESIS as described by Song et al. (*45*). The p-values reported for our logistic regression analysis were calculated using the saddle point approximation (*45*). We, again, used the same GRM and PCs as described in section 5.1, and adjusted by the same covariates as our linear regression analysis, without the batch variable. We used the Bonferroni method to adjust for multiple hypothesis testing. However, we adjusted each analysis separately and only by the number of transcripts tested in that analysis. Any transcripts for which the GENESIS logistic regression model failed to converge were excluded from analysis. Thus, for the logistic regression, we only analyzed 846 transcripts.

#### 5.3.2 Gene Set Enrichment Analyses

We performed gene set enrichment analyses using the hypeR package (*55*). We tested for enrichment in the Kyoto Encyclopedia of Genes and Genomes (KEGG) and hallmark compendiums downloaded from MsigDB. We tested using two different backgrounds: the overlap between the gene set genes and all the tested transcripts and the overlap between the gene set genes and all the tested logistic transcripts (transcripts in the logistic regression data set). We tested signatures from the linear transcripts that reached nominal significance, the logistic transcripts that reached nominal significance and the combined transcript set of the linear and logistic nominally significant transcripts. We further divided and tested the linear, logistic and combined signatures based on the domain of the associated SNPs, the test of the associated SNPs and the associated SNP. We excluded transcripts without a valid HGNC symbol from analysis. We define gene sets as enriched based on q < 0.05 using the Benjamini-Hochberg correction method.

#### 5.3.3 Metabolite QTLs (mQTLs)

We performed linear regression analyses to identify associations between metabolites and our genome-wide significant SNPs. We used the same QTL pipeline, GRM and PCs as described in section 5.1. We adjusted by sex, field center, education, birth cohort, the participants age at the time of the blood draw, the interaction between age and age cohort, and indicator variables for hypertension, type 2 diabetes, lipid lowering, and heart medication use. We used the Bonferroni method to adjust for multiple hypothesis testing.

We define nominal significance as p<0.05 (unadjusted). We treated polar metabolites and lipid metabolites as separate analyses and determined the Bonferroni adjusted significance according to the number of metabolites available in that analysis.

## Figure Captions

**Supplementary Figure 1:**
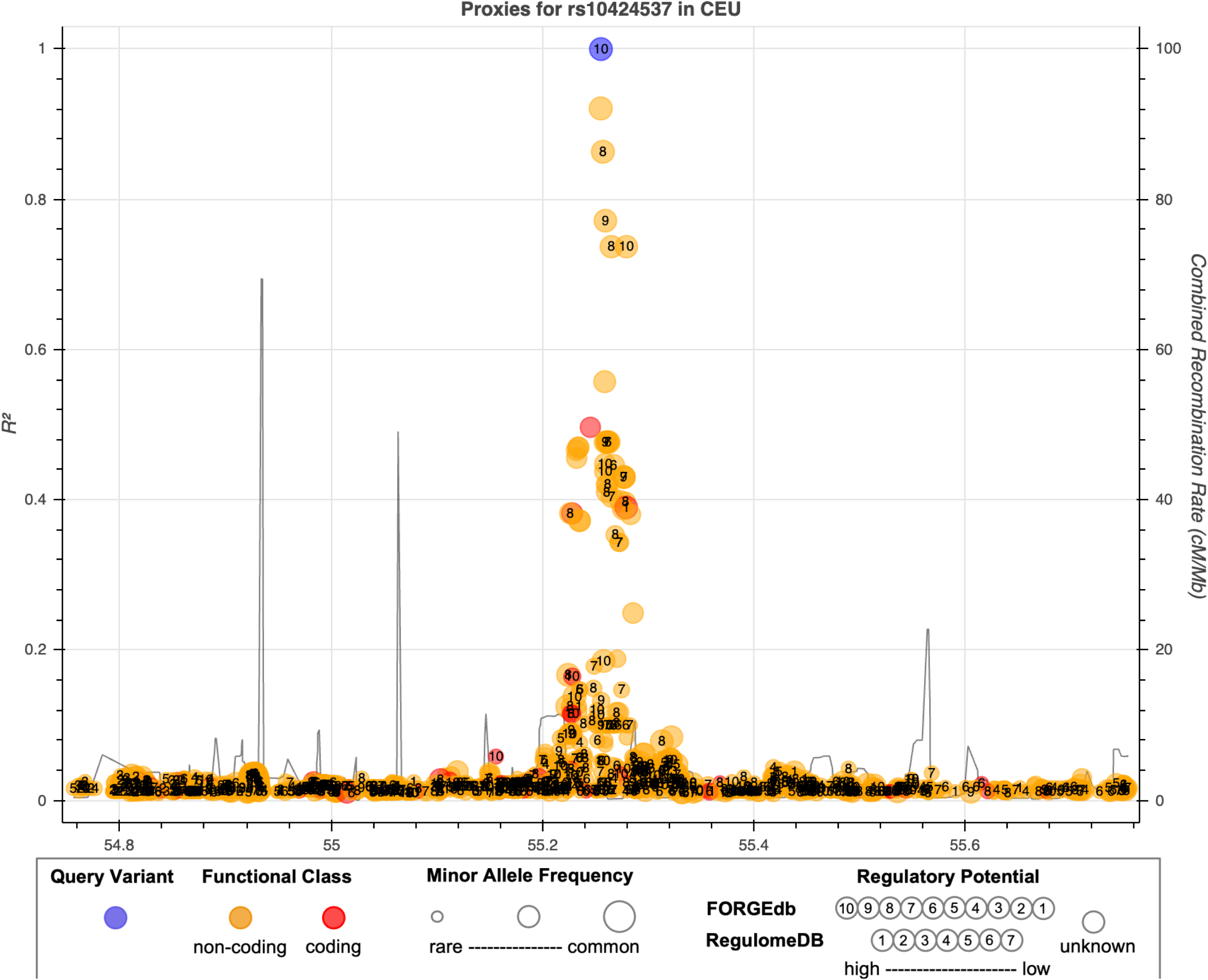
Ldlink Brokeh plot of SNPs proximal to rs10424537. Each point indicates the correlation between rs10424537 and another point within ±500kb. The number within each point is the FORGEdb score. The point size indicates the position’s minor allele frequency. Each point is color coded according to the whether the positi on is the querry variant (rs10424537, blue), coding (exonic, red) or non-coding (everything else, yellow). rs4473307 (*R*^2^ = 0.921) and rs12983085 (*R*^2^ = 0.8633) are the only positions with correlations above 0.8, and they are just under genome wide significance in our GWAS data (see Supplementary table 13).

**Supplementary Figure 2:**
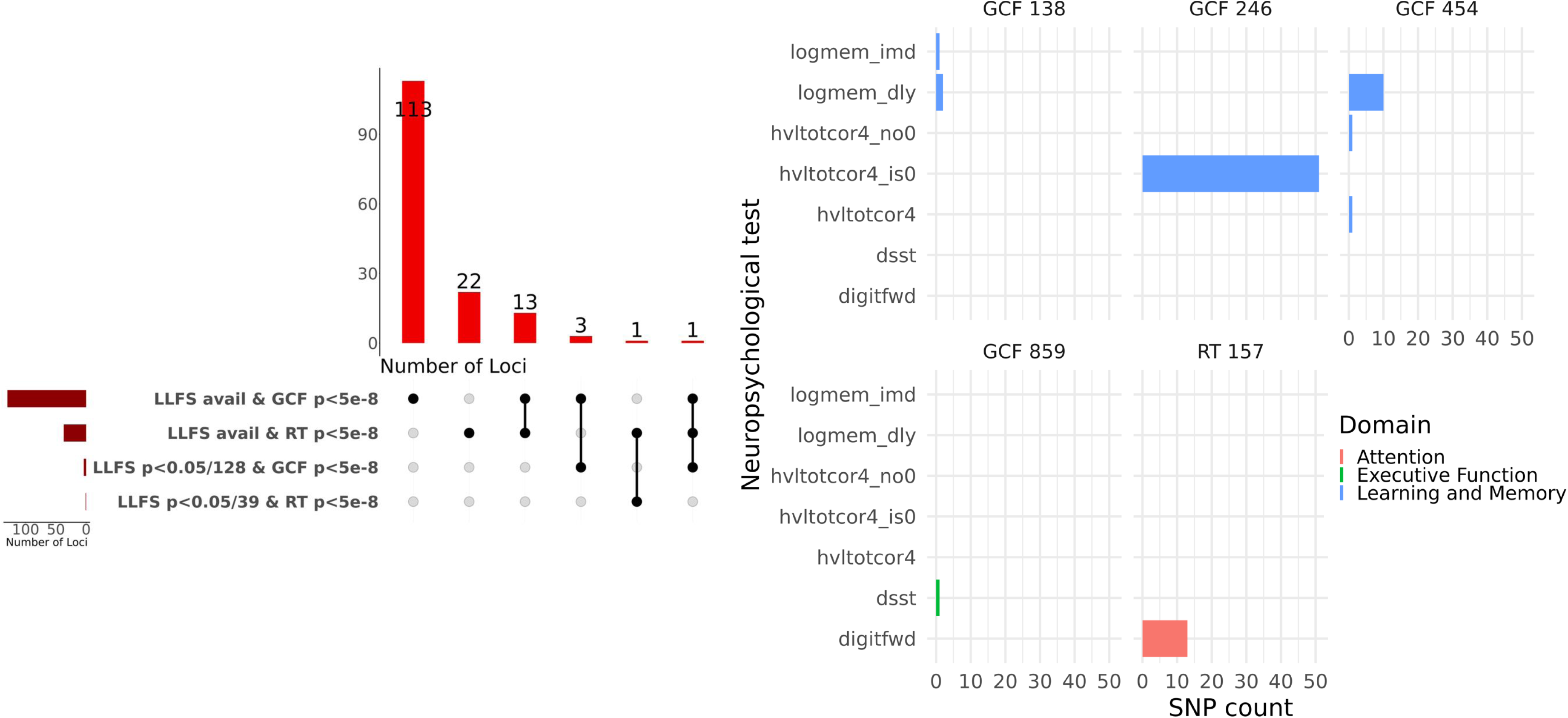
Comparison of Davies et. al Results by Loci and Domain. **Figure A:** Upset plot of the loci available in the LLFS and Davies et. al GWASs broken down by significance. **Figure B:** Bar Plots of the number of SNPs associated in each test that come from a given locus color coded by the domain of the test. See methods section 5.2 for more details.

**Supplementary Figure 3:**
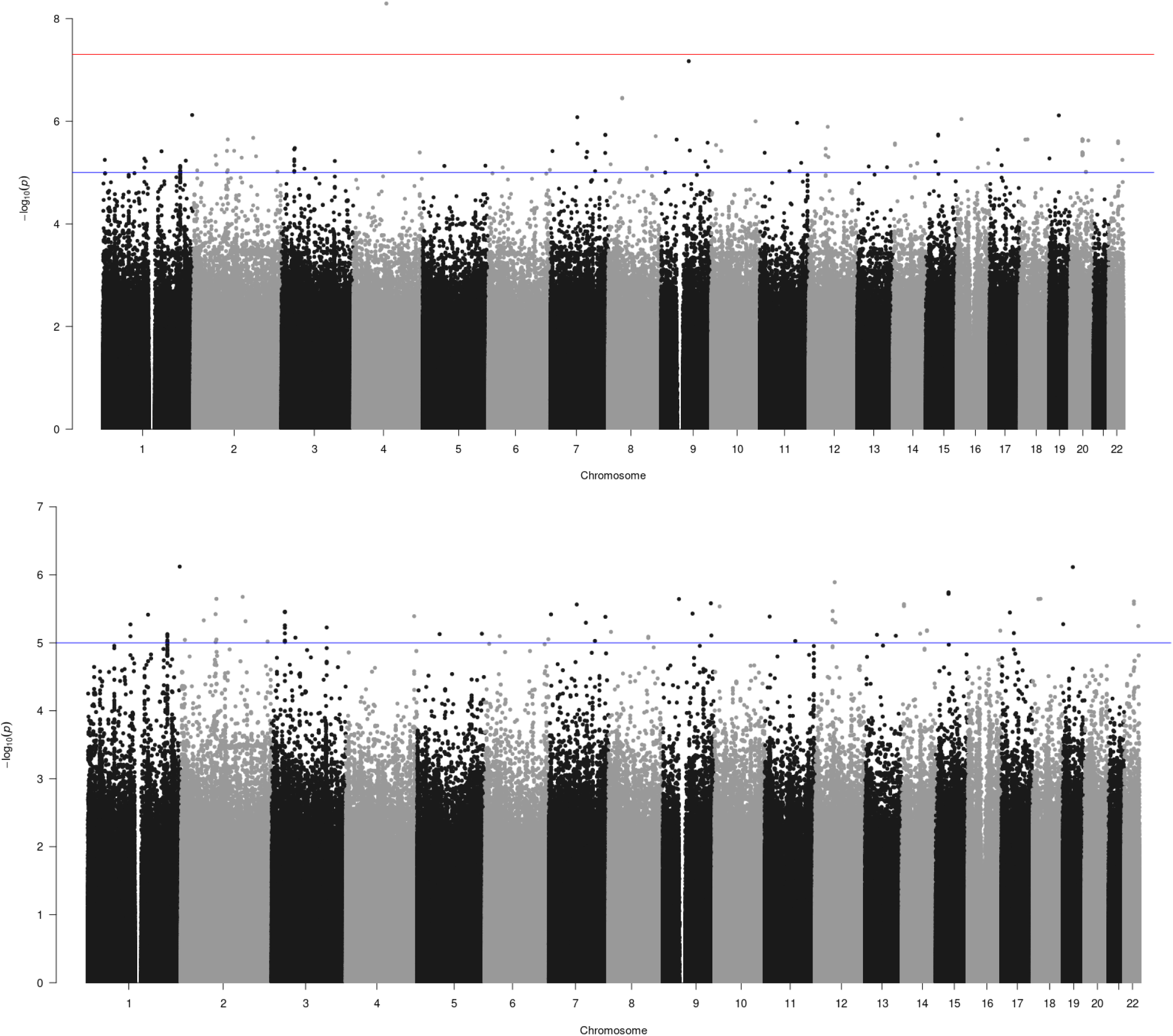
Semantic Fluency Test Manhattan Plots. Mac 3 on Top and Mac 5 on bottom.

**Supplementary Figure 4:**
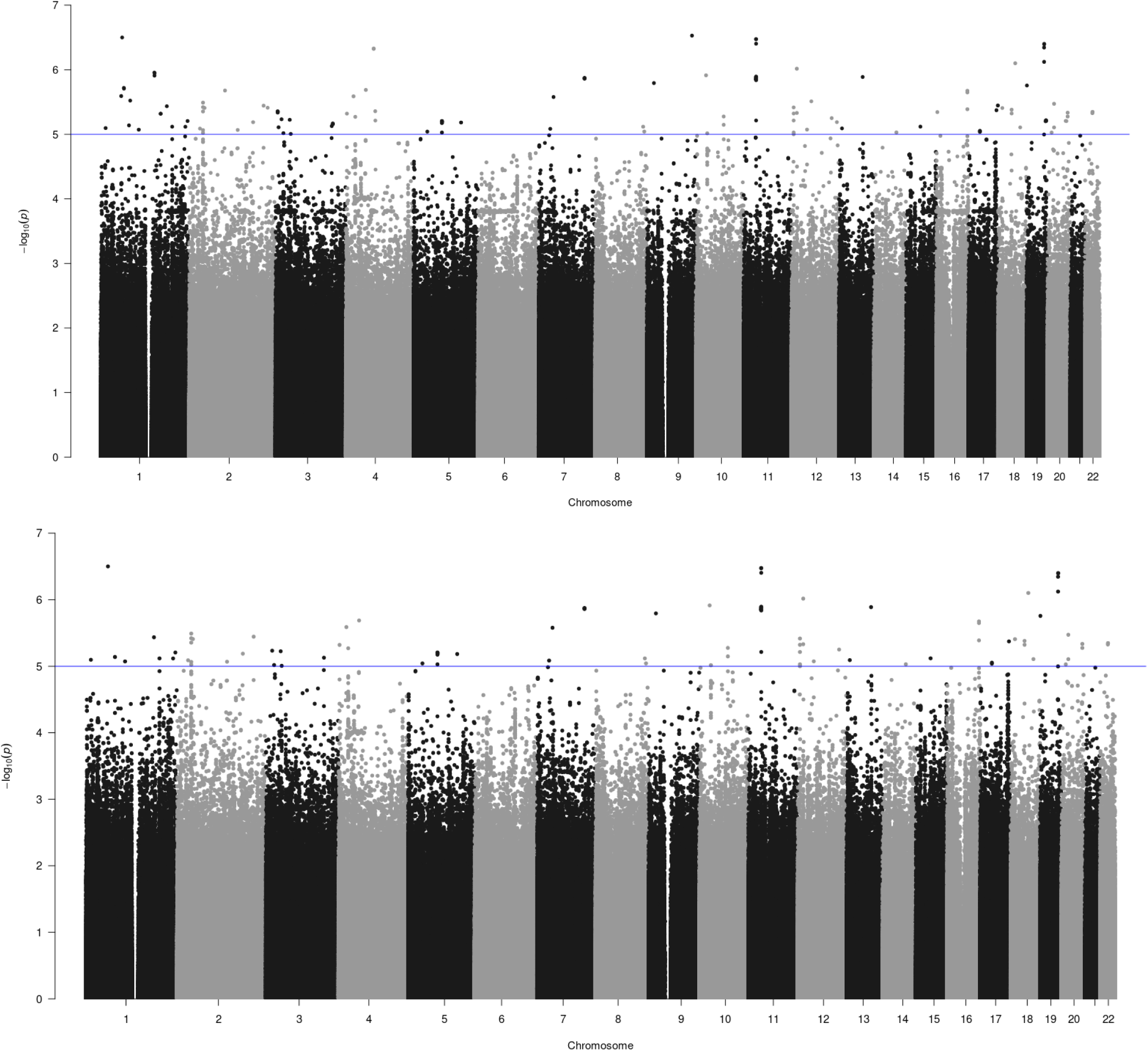
Number Span Backward Manhattan Plots. Mac 3 on Top and Mac 5 on bottom.

**Supplementary Figure 5:**
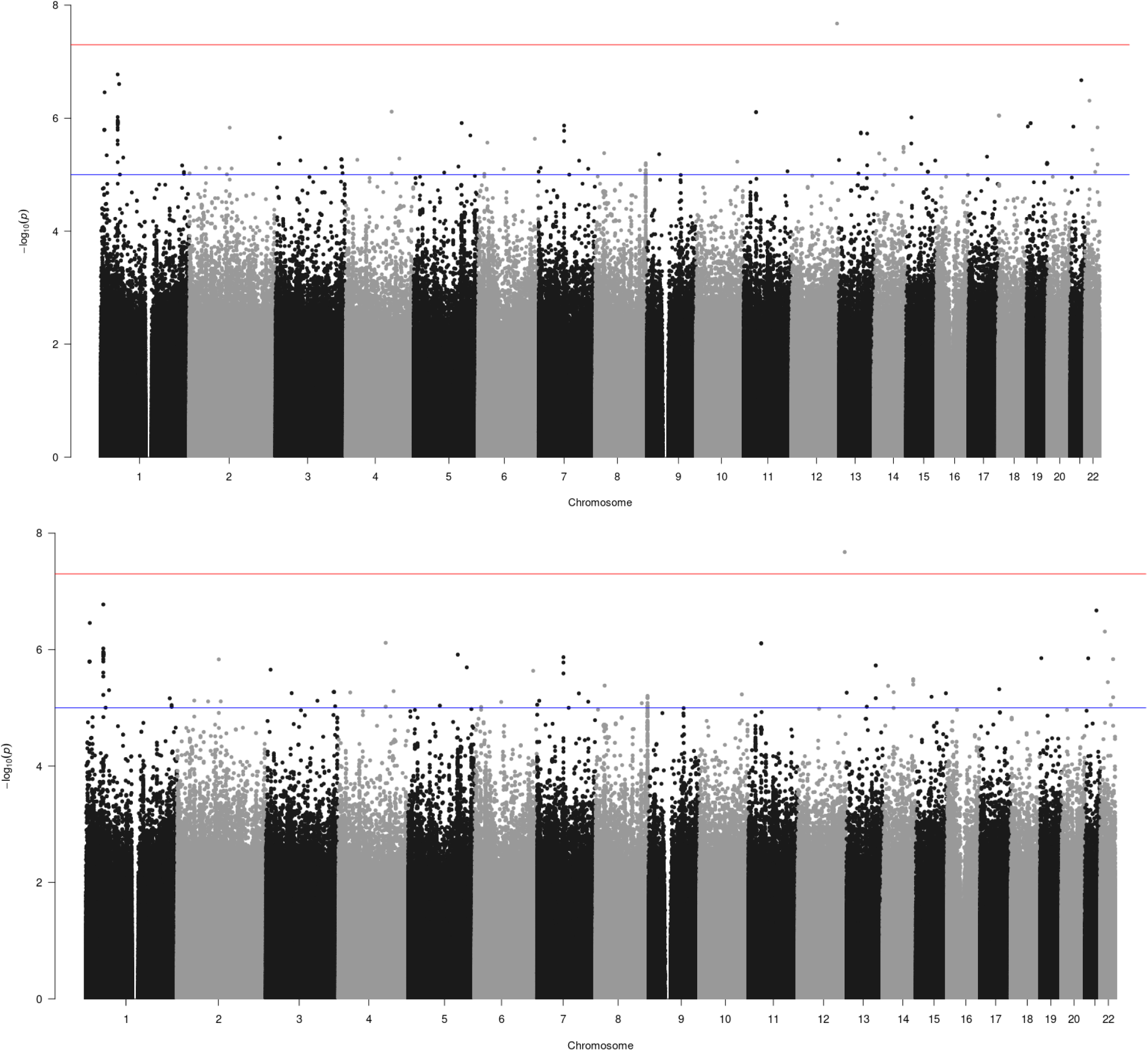
Number Span Forward Manhattan Plots. Mac 3 on Top and Mac 5 on bottom.

**Supplementary Figure 6:**
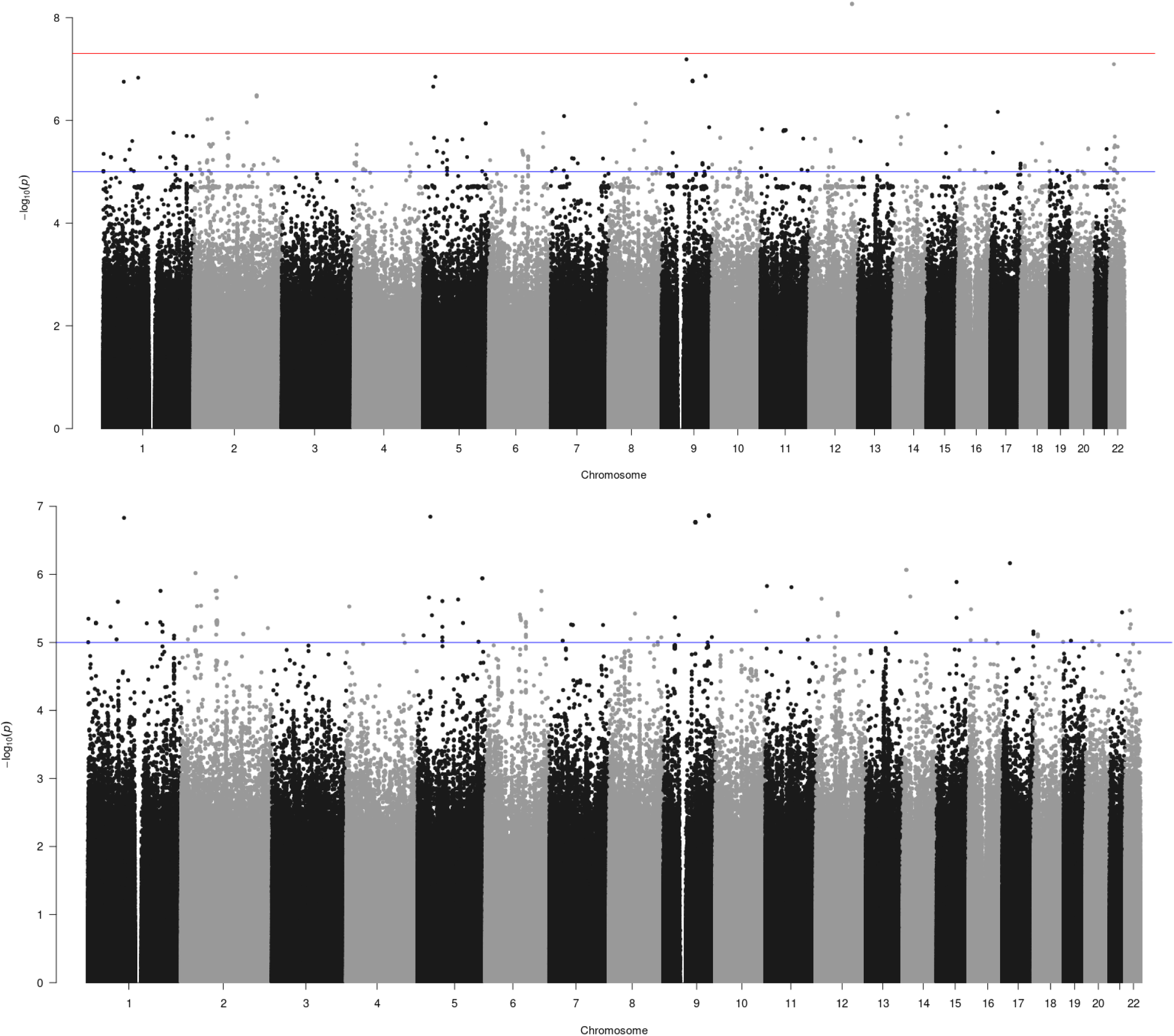
Digit Symbol Substitution Test Manhattan Plots. Mac 3 on Top and Mac 5 on bottom.

**Supplementary Figure 7:**
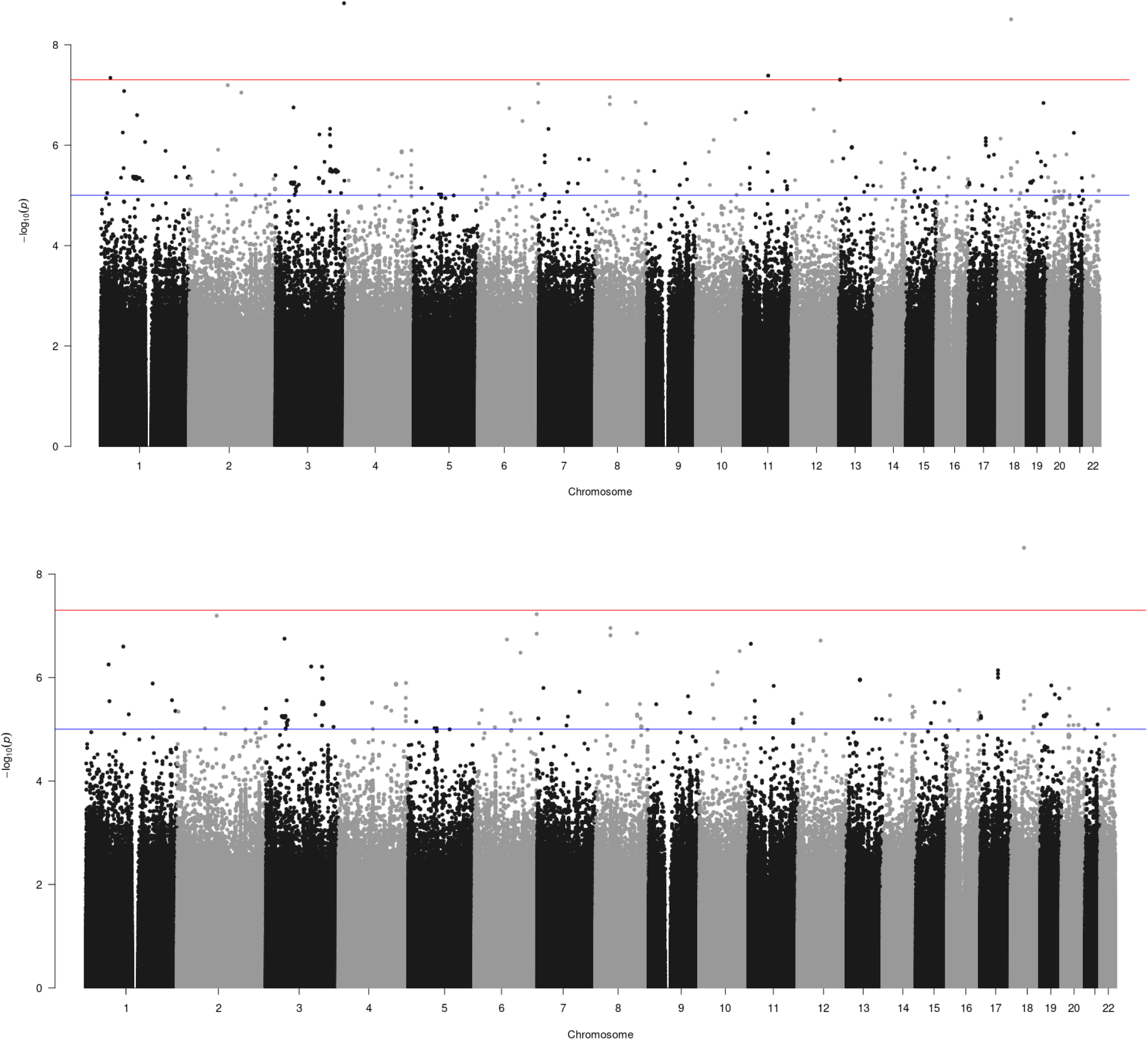
Controlled Oral Word Association Test Manhattan Plots. Mac 3 on Top and Mac 5 on bottom.

**Supplementary Figure 8:**
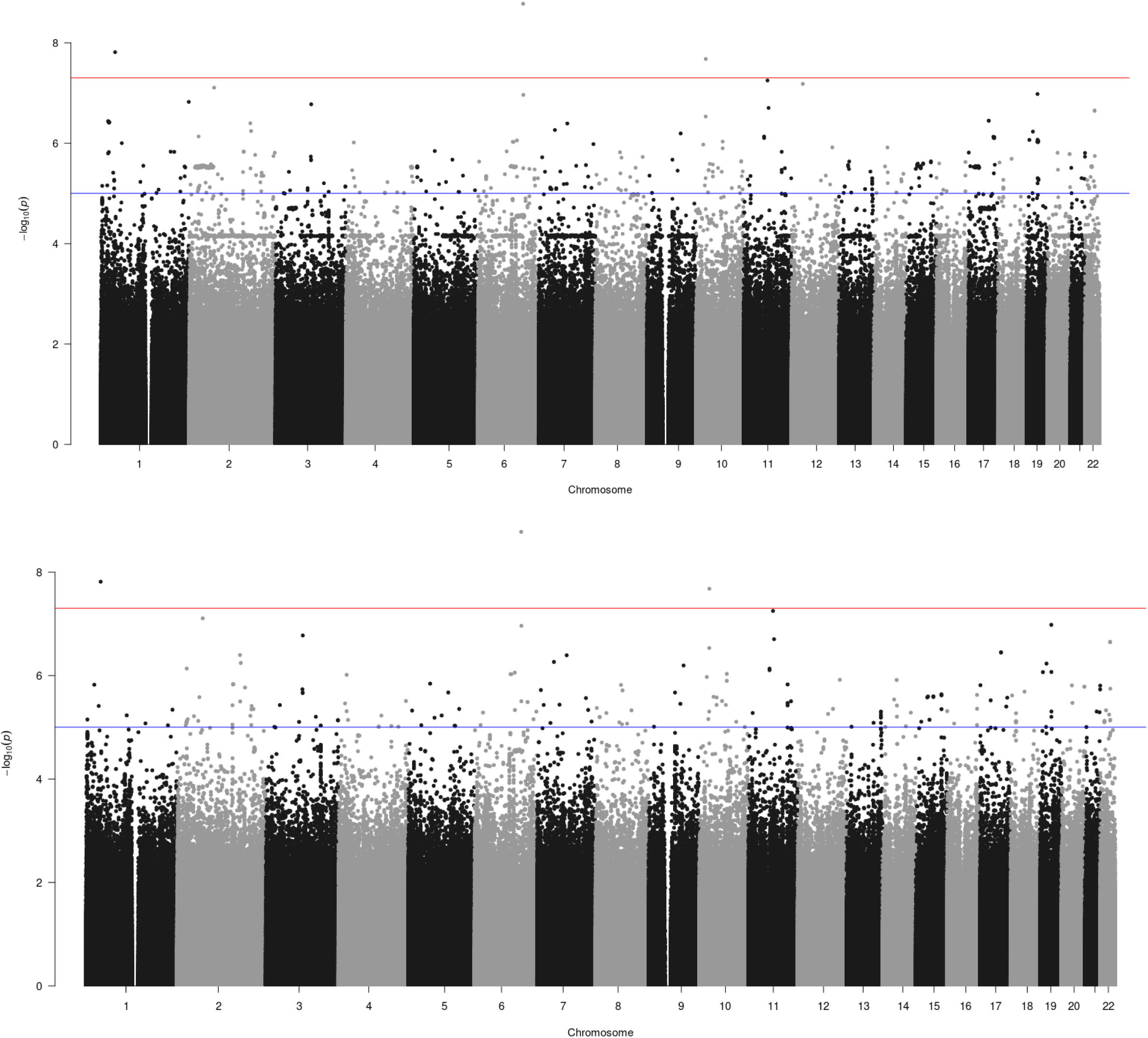
Hopkins Verbal Learning Test –Revised delayed recall Manhattan Plots. Mac 3 on Top and Mac 5 on bottom.

**Supplementary Figure 9:**
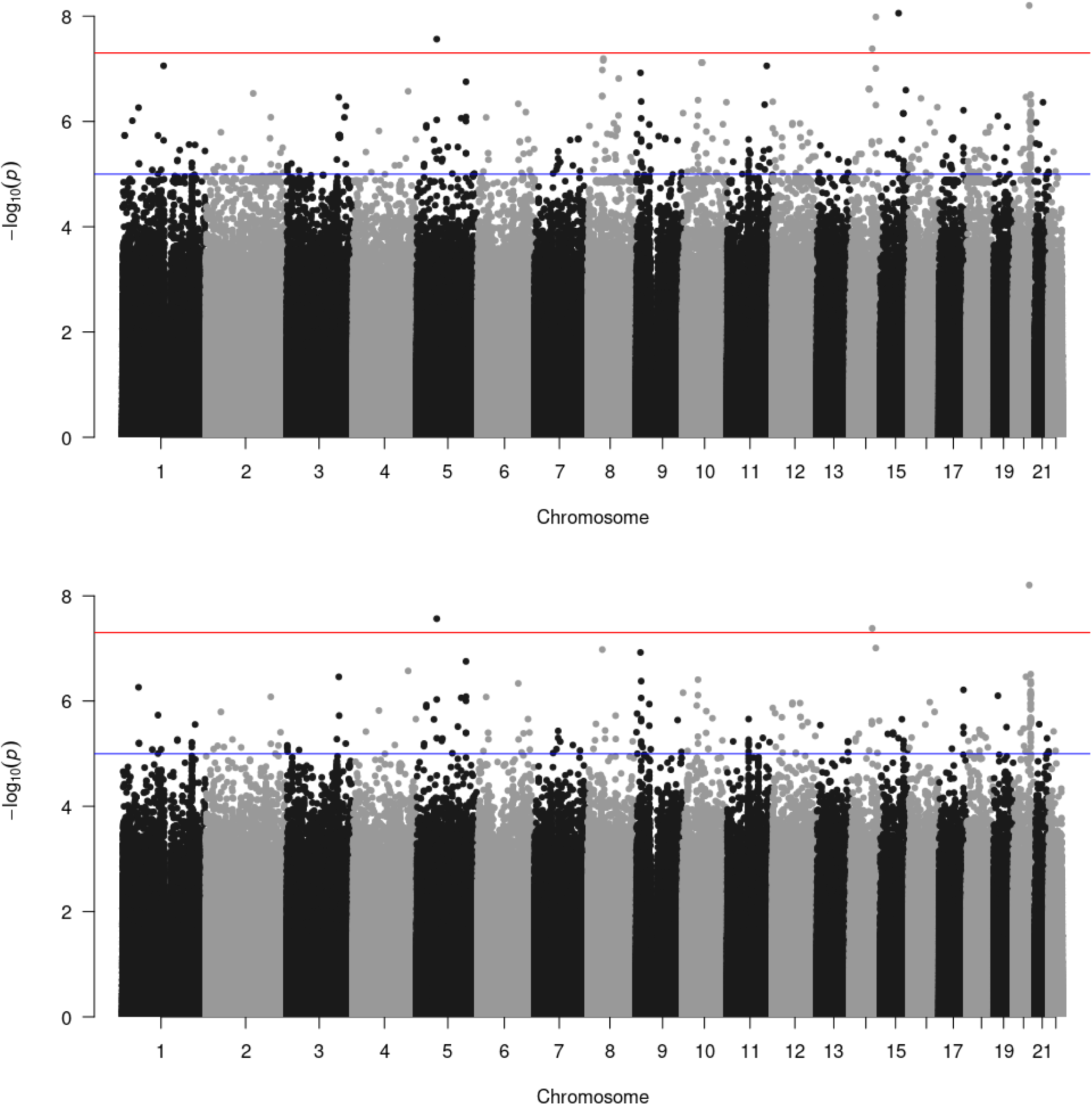
Hopkins Verbal Learning Test –Revised delayed recall, samples scoring 0 excluded Manhattan Plots. Mac 3 on Top and Mac 5 on bottom.

**Supplementary Figure 10:**
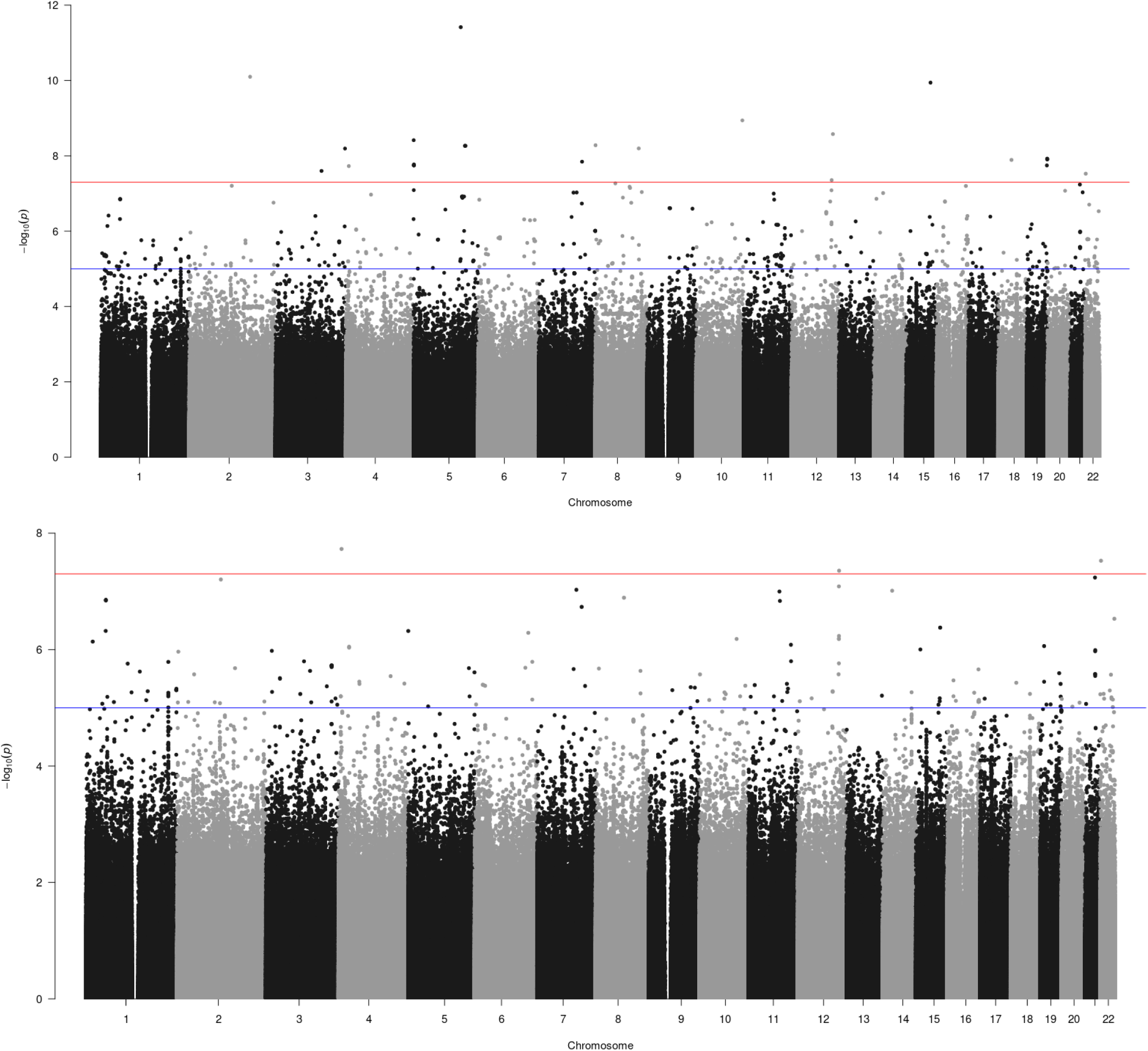
Hopkins Verbal Learning Test –Revised learning trials Manhattan Plots. Mac 3 on Top and Mac 5 on bottom.

**Supplementary Figure 11:**
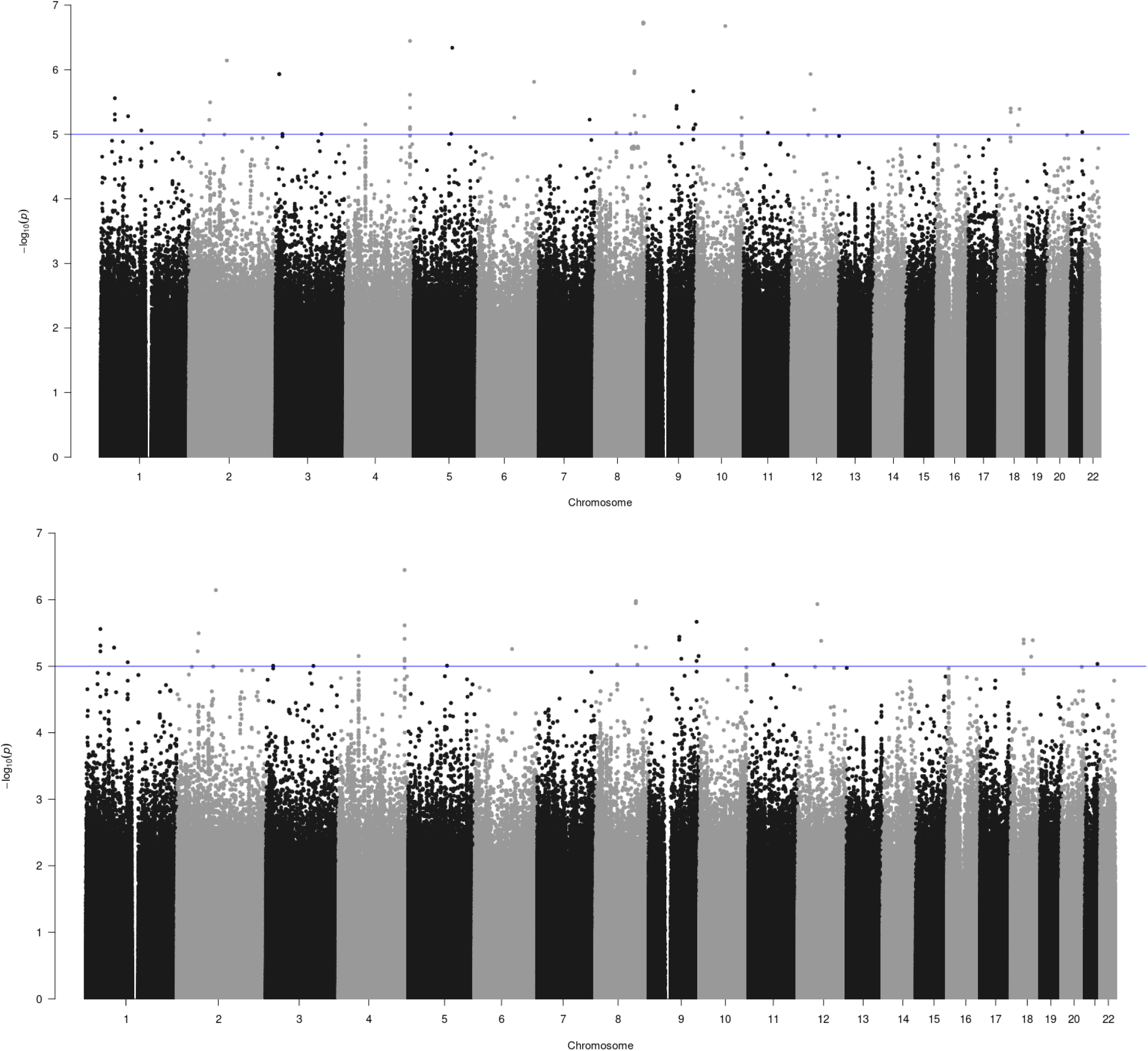
Logical Memory IIA from the Wechsler Memory Scale – Revised Manhattan Plots. Mac 3 on Top and Mac 5 on bottom.

**Supplementary Figure 12:**
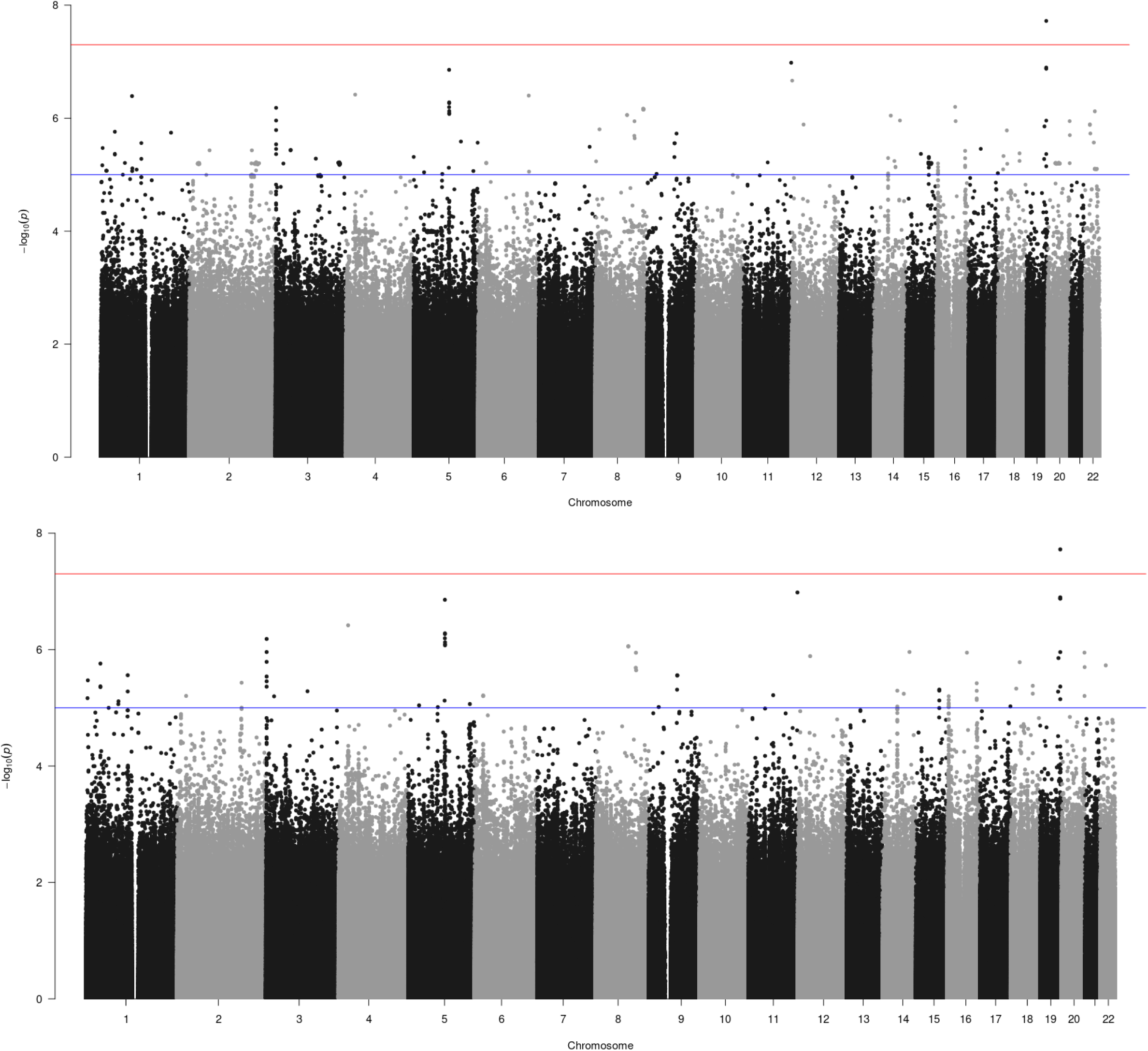
Logical Memory IA from the Wechsler Memory Scale – Revised Manhattan Plots. Mac 3 on Top and Mac 5 on bottom.

## Notes

### Competing Interest Statement

The authors have declared no competing interest.

